# Chromatin and transcriptional response to loss of TBX1 in early differentiation of mouse cells

**DOI:** 10.1101/2020.06.06.137026

**Authors:** Andrea Cirino, Ilaria Aurigemma, Monica Franzese, Gabriella Lania, Dario Righelli, Rosa Ferrentino, Elizabeth Illingworth, Claudia Angelini, Antonio Baldini

## Abstract

The T-box transcription factor TBX1 has critical roles in the cardiopharyngeal lineage and the gene is haploinsufficient in DiGeorge syndrome, a typical developmental anomaly of the pharyngeal apparatus. Despite almost two decades of research, if and how TBX1 function triggers chromatin remodeling is not known.

Here, we explored genome-wide gene expression and chromatin remodeling in two independent cellular models of *Tbx1* loss of function, mouse embryonic carcinoma cells P19Cl6, and mouse embryonic stem cells (mESCs). The results of our study revealed that the loss or knockdown of TBX1 caused extensive transcriptional changes, some of which were cell type-specific, some were in common between the two models. However, unexpectedly we observed only limited chromatin changes in both systems. In P19Cl6 cells, differentially accessible regions (DARs) were not enriched in T-BOX binding motifs; in contrast, in mESCs, 34% (n=47) of all DARs included a T-BOX binding motif and almost all of them gained accessibility in *Tbx1*^-/-^ cells.

In conclusion, despite a clear transcriptional response of our cell models to loss of TBX1 in early cell differentiation, chromatin changes were relatively modest.

## INTRODUCTION

TBX1 is a transcription factor encoded by a gene that is haploinsufficient in DiGeorge / 22q11.2 deletion syndromes and in the mouse (Baldini et al., 2017; Greulich et al., 2011; McDonald-McGinn et al., 2015). It is a critical player in the development of the pharyngeal apparatus, which gives rise to organs and structures that are affected by many birth defects. The mechanisms by which TBX1 regulates transcription are only now beginning to emerge, but many questions remain. It binds DNA to a typical T-BOX consensus motif (Castellanos et al., 2014; Fulcoli et al., 2016), and interacts with transcription regulators such as chromatin remodeling complexes, histone modifiers, as well as repressive co-factors (Chen et al., 2012; Fulcoli et al., 2016; Okubo et al., 2011; Stoller et al., 2010), and positively regulates H3K4 monomethylation (Fulcoli et al., 2016). In the mouse, *Tbx1* is expressed early in development (from around E7.5) in the cardiopharyngeal mesoderm, the developing anterior foregut/pharyngeal endoderm and in the surface ectoderm. Timed-deletion of the gene has revealed a requirement as early as E7.5-E8.0 (Xu et al., 2005) for the development of the 4th pharyngeal arch artery that will form much later. While this phenomenon could be explained by a number of mechanisms, one possibility is that TBX1 primes enhancers for downstream activation or repression, thereby creating asynchrony between the time of requirement and the phenotypic consequences. To address this issue, we have used two cellular models that respond transcriptionally to *Tbx1* gene dosage, mouse P19Cl6 and embryonic stem cells (mESCs), but that are at an early stage of differentiation, and we tested the effects of *Tbx1* inactivation on transcription and on chromatin remodeling. mESCs (*Tbx1*^+/+^ and *Tbx1*^-/-^) were subjected to a widely used cardiac mesoderm differentiation protocol (Kattman et al., 2011) and selected using a fluorescence activated cell sorting (FACS) approach. We selected a subpopulation that expresses the highest level of *Tbx1* and pursued it for ATAC-seq (Buenrostro et al., 2015) and RNA-seq. The results obtained from the two cellular models indicate that TBX1 inactivation does not have a strong effect on chromatin remodeling at the differentiation stages tested despite having significant effects on transcription. We discuss possible mechanisms to explain our results.

## MATERIALS AND METHODS

### P19Cl6 cells

We plated 5×10^5^ cells in a 35-mm dish in Dulbecco-Modified Minimal Essential Medium supplemented (Sigma-Aldrich #M4526) with 10% fetal bovine serum (Gibco #10270106). After 24 hours, at confluence, we added 10uM 5-Azacytidine (5-Aza, Sigma-Aldrich #A2385) to induce differentiation. For ATAC-seq, cells were harvested after a further 24 hr. For quantitative ATAC experiments in time course, 24 hr after 5-Aza treatment we replaced the media with fresh media containing 1% DMSO. Samples for qATAC were harvested 13 hr after transfection (T1), 24 hr after 5-Aza induction (D1), and 24 hr after DMSO treatment (D2).

### Mouse embryonic stem cells (mESCs)

E14-Tg2a mESCs were cultured without feeders and maintained undifferentiated on gelatin-coated dishes in GMEM (Sigma Cat# G5154) supplemented with 10^3^U/ml ESGRO LIF (Millipore, Cat# ESG1107), 15% fetal bovine serum (ES Screened Fetal Bovine Serum, US Euroclone Cat# CHA30070L), 0.1 mM nonessential amino acids (Gibco, Cat# 11140-035), 0.1 mM 2-mercaptoethanol (Gibco, Cat# 31350-010), 0.1 mM L-glutamine (Gibco, Cat# 25030081), 0.1 mM Penicillin/Streptomycin (Gibco, Cat# 10378016), and 0.1 mM sodium pyruvate (Gibco, Cat# 11360-070). The cells were passaged every 2-3 days using 0.25% Trypsin-EDTA (1X) (Gibco, Cat# 25200056) as the dissociation buffer.

For differentiation, E14-Tg2a mESCs were dissociated with Trypsin-EDTA and cultured at 75,000 cells/ml in serum-free medium: 75% Iscove’s modified Dulbecco’s medium (Cellgro Cat# 15-016-CV) and 25% HAM F12 media (Cellgro #10-080-CV), supplemented with N2 (GIBCO #17502048) and B27 (GIBCO #12587010) supplements, penicillin/streptomycin (GIBCO #10378016), 0.05% BSA (Invitrogen Cat#. P2489), L-glutamine (GIBCO #25030081), 5mg/ml ascorbic acid (Sigma A4544) and 4.5 × 10^−4^ M monothioglycerol (Sigma M-6145). After 48 hr in culture, the EBs were dissociated with trypsin-EDTA and reaggregated for 40 hr in serum-free differentiation media with the addition of 8ng/ml human activin A (R&D Systems Cat#. 338-AC), 0.5 ng/ml human BMP4 (R & D Systems Cat# 314-BP), and 5ng/ml human VEGF (R&D Systems Cat#. 293-VE). The 2-day-old EBs were dissociated and 6×10^4^ cells were seeded onto individual wells of a 24-well plate coated with 0.1% gelatin in StemPro-34 medium (Gibco #10639011), supplemented with SP34 supplement, L-glutamine, 5mg/ml ascorbic acid, 5 ng/ml human-VEGF, 10 ng/ml human bFGF (R&D Systems 233-FB-025), and 50 ng/ml human FGF10 (R&D Systems 338-FG-025). After 48 hr, we added new StemPro-34 media, supplemented with SP34 supplement, L-glutamine, 5mg/ml ascorbic acid and keep for 96 hr. We harvested cells for RNA-seq and ATAC-seq analysis at day 4 of differentiation (Fig. 1A).

**Figure 1.**
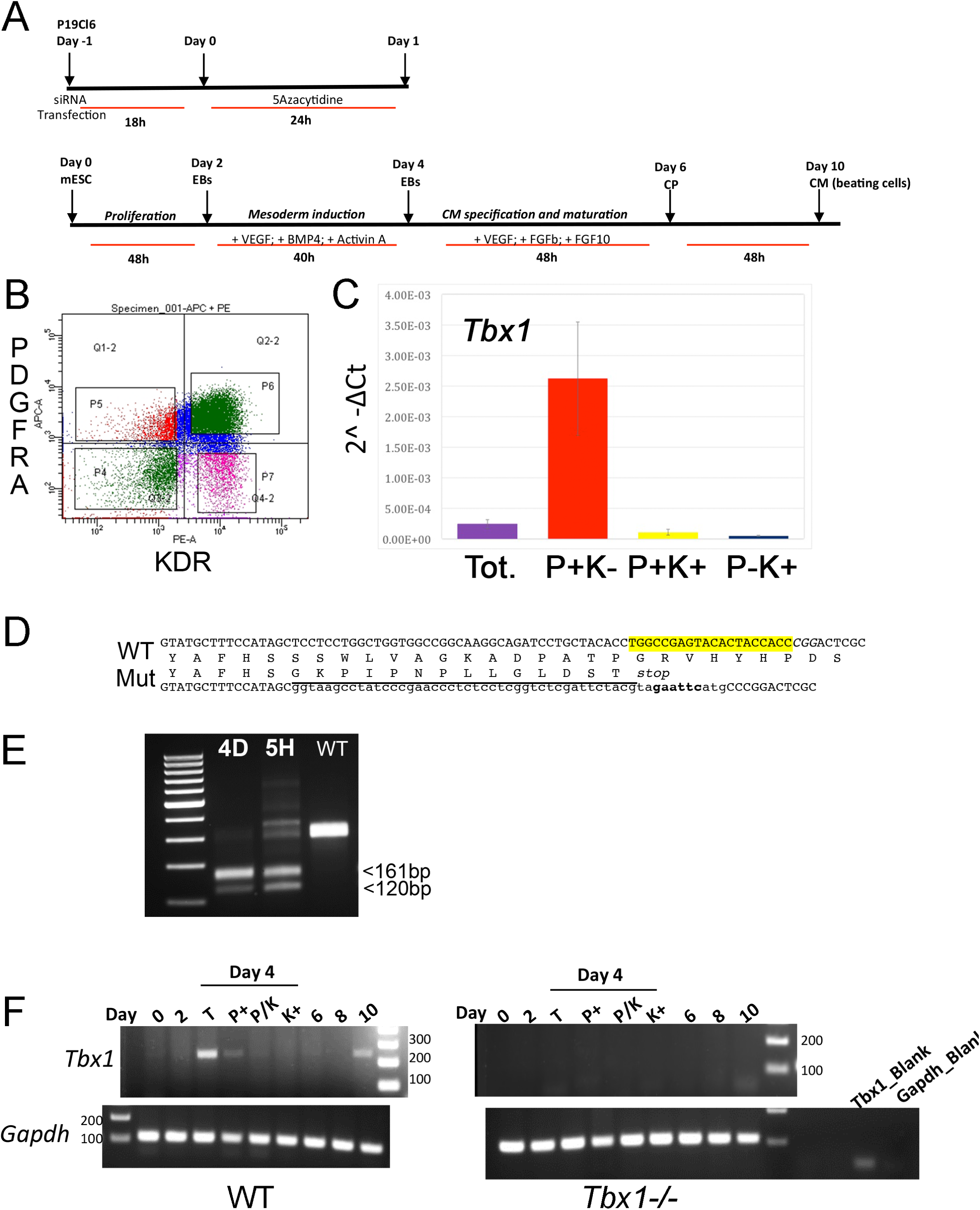
Experimental protocols and reagents used in this study. A) Top: differentiation protocols for P19Cl6 cells, transfected with *Tbx1*-targeted or non-targeted siRNA. Cells were assayed at day 1 of differentiation. Bottom: differentiation protocol used for mES cells. Cells were assayed at day 4). B). Representative plot of the gating strategy used for immunophenotyping of cell during mES differentiation. The PDGFRA+;KDR-, PDGFRA+;KDR+, PDGFRA-;KDR+ subpopulations were identified at day 4 of differentiation by FACS using anti-PGFRA and anti-KDR antibodies. C) Quantitative real time PCR. *Tbx1* expression was evaluated in the three subpopulations and in unsorted cells (Tot.). Abbreviations: P, PDGFRA ; K, KDR. D) Strategy to generate a knockout allele of *Tbx1* using CRISPR-Cas9 and homologous recombination. The top line shows the WT sequence of ex 3/7 (or 5/9). In yellow, the gRNA sequence; in italic the PAM sequence, CGG. The two intermediate lines indicate the predicted amino acid sequences. The bottom line indicates the sequence of the recombinant allele. WT sequence is shown in uppercase; the sequence inserted by homologous recombination is shown in lowercase, this includes a V5 tag (underlined), a stop codon, and a diagnostic *Eco*RI digestion site (in bold). E) PCR amplification of the targeted region from *Tbx1* homozygous clones 4D and 5H, and from WT digested with *Eco*RI. F) *Tbx1* expression revealed by reverse transcription PCR. Left panel : PCR of samples collected at the differentiation stages indicated on WT mES cells. At day 4, the analysis was performed on total populations (T) and on FACS-purified subpopulations. The right panel shows the same experiment performed using the *Tbx1*^*-/-*^ clone 5H. Abbreviations: P, PDGFRA ; K, KDR.

### CRISPR-Cas9-mediated targeting of mESCs

*Tbx1* knockout was induced in E14-Tg2a using Alt-R™ CRISPR-Cas9 System (IDT) following the manufacturer’s specifications. This genome editing system is based on the use of a ribonucleoprotein (RNP) consisting of Alt-R S.p. Cas9 nuclease complexed with an Alt-R CRISPR-Cas9 guide RNA (crRNA:tracrRNA duplex). The crRNA is a custom synthesized sequence that is specific for the target (Tbx1KO: /AltR1/rUrG rGrCrC rGrArG rUrArC rArCrU rArCrC rArCrC rGrUrU rUrUrA rGrArG rCrUrA rUrGrC rU/AltR2/) and contains a 16 nt sequence that is complementary to the tracrRNA. Alt-R CRISPR-Cas9 tracrRNA-ATTO 550 (5nmol catalog n. 1075927) is a conserved 67 nt RNA sequence that is required for complexing to the crRNA so as to form the guide RNA that is recognized by S.p. Cas9 (Alt-R S.p. Cas9 Nuclease 3NLS, 100 µg catalog n. 1081058). The fluorescently labeled tracrRNA with ATTO™ 550 fluorescent dye is used to FACS-purify transfected cells. The protocol involves 3 steps: 1) annealing of the crRNA and tracrRNA, 2) assembly of the Cas9 protein with the annealed crRNA and tracrRNAs, and 3) delivery of the ribonucleoprotein (RNP) complex into mESC by reverse transfection. Briefly, we annealed equimolar amounts of resuspended crRNA and tracrRNA to a final concentration (duplex) of 1 μM by heating at 95°C for 5 min and then cooling to room temperature. The RNA duplexes were then complexed with Alt-R S.p. Cas9 enzyme in OptiMEM medium to form the RNP complex, which was then transfected into mESCs using the RNAiMAX transfection reagent (Invitrogen). After 48 hr incubation, cells were trypsinized and ATTO 550+ (transfected) cells were purified by FACS. Fluorescent cells (approx. 65% of the total cell population) were plated at very low density to facilitate colony picking. We picked and screened by PCR 96 clones. Positive clones were confirmed by DNA sequencing.

### Fluorescence activated cell sorting (FACS)

For FACS, EBs were collected and allowed to settle by gravity. After washing with PBS, the cells were dissociated using the Embryoid Body dissociation kit (cod. 130-096-348 Miltenyi Biotec) according to the manufacturer’s protocol. Dissociated cells (1x 10^6^ cells/100 μl) were incubated with primary antibodies (PDGFRα-APC, mouse cod.130-102-473; KDR VEGFR2-PE (KDR), mouse cod. 130-120-813 Miltenyi Biotec) directly conjugated (1:50) in PBS-BE solution (PBS, 0.5%BSA, 5mM EDTA) for 20 min on ice. Subsequently, cells were washed twice with 2ml of PBS-BE. Cells were sorted using the BD FACS ARIAIII™ cell sorter.

Total RNA was isolated using QIAzol lysis reagent (QIAGEN) and for qRT-PCR it was reverse-transcribed using the High Capacity cDNA Reverse Transcription kit (Applied Biosystem catalog. n. 4368814). Quantitative real-time PCR was performed using SYBR Green PCR master mix (Applied Biosystem catalog. n. 4309155). Relative gene expression was evaluated using the 2^-ΔCt method, and *Gapdh* expression as the normalizer. Primer sequences are listed on Tab. 2.

**Table 1.**
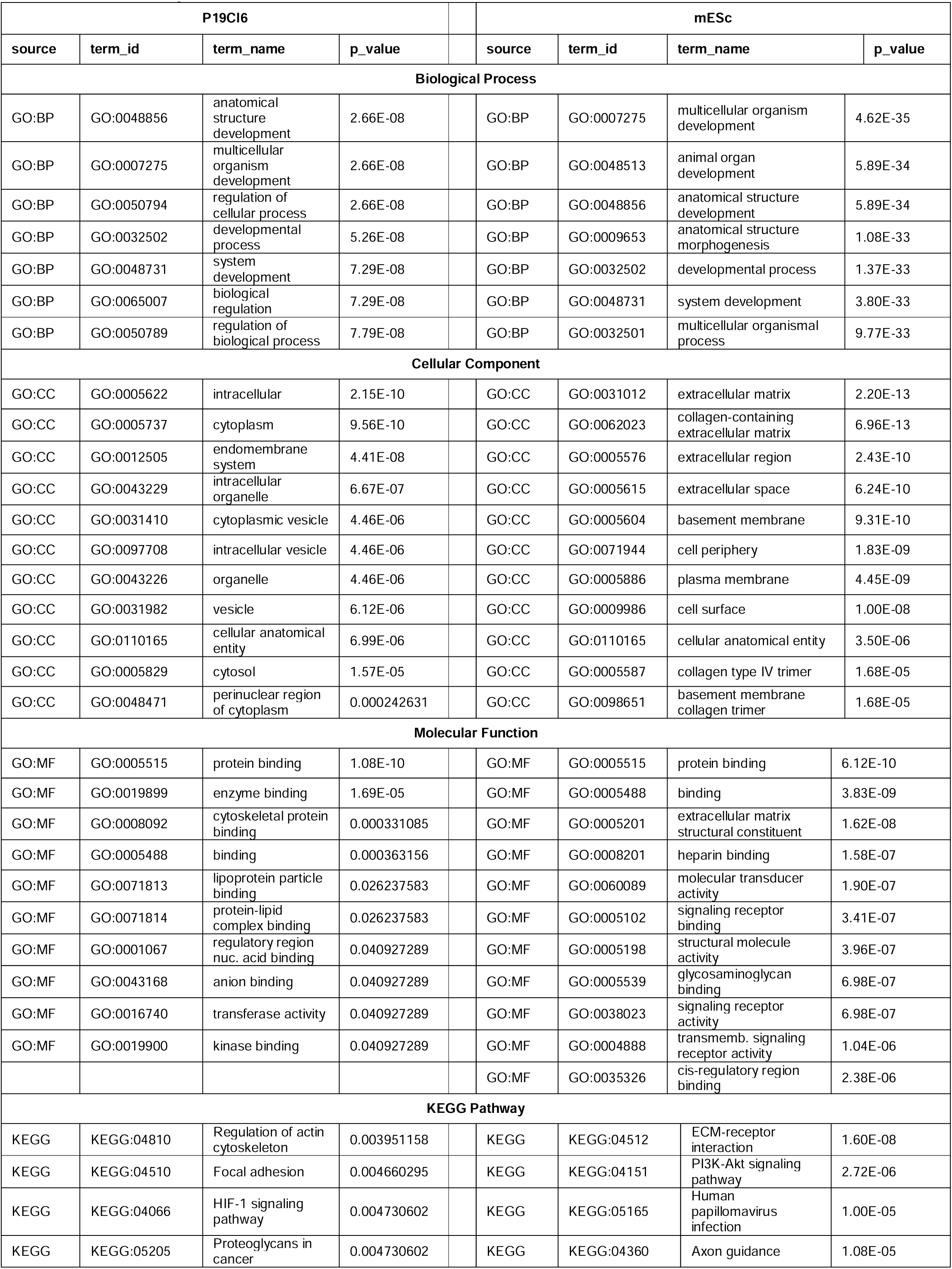
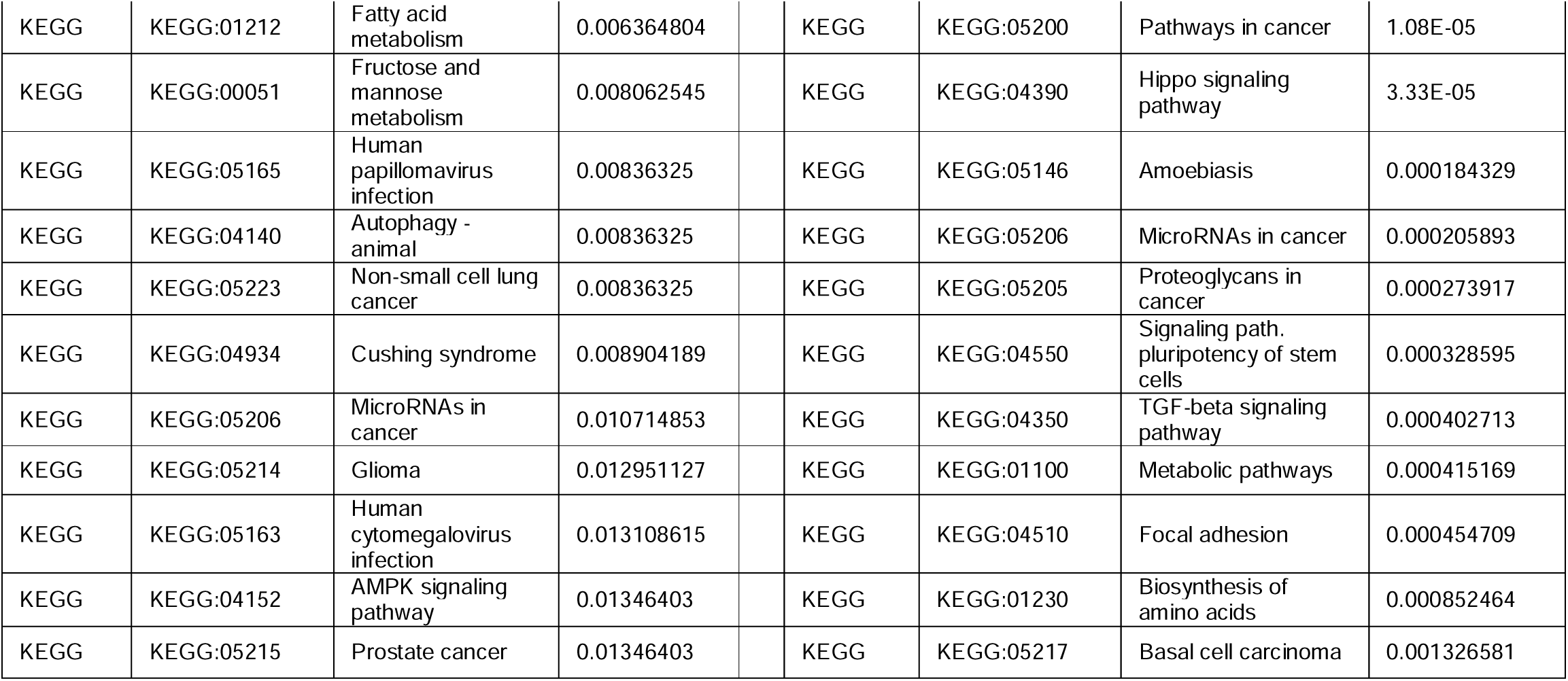
(GO analysis)

| P19C16 |  |  |  | mESc |  |  |  |
| --- | --- | --- | --- | --- | --- | --- | --- |
| source | term_id | term_name | p_value | source | term_id | term_name | p_value |
| **Biological Process** |  |  |  |  |  |  |  |
| GO:BP | GO:0048856 | anatomical structure development | 2.66E-08 | GO:BP | GO:0007275 | multicellular organism development | 4.62E-35 |
| GO:BP | GO:0007275 | multicellular organism development | 2.66E-08 | GO:BP | GO:0048513 | animal organ development | 5.89E-34 |
| GO:BP | GO:0050794 | regulation of cellular process | 2.66E-08 | GO:BP | GO:0048856 | anatomical structure development | 5.89E-34 |
| GO:BP | GO:0032502 | developmental process | 5.26E-08 | GO:BP | GO:0009653 | anatomical structure morphogenesis | 1.08E-33 |
| GO:BP | GO:0048731 | system development | 7.29E-08 | GO:BP | GO:0032502 | developmental process | 1.37E-33 |
| GO:BP | GO:0065007 | biological regulation | 7.29E-08 | GO:BP | GO:0048731 | system development | 3.80E-33 |
| GO:BP | GO:0050789 | regulation of biological process | 7.79E-08 | GO:BP | GO:0032501 | multicellular organismal process | 9.77E-33 |
| **Cellular Component** |  |  |  |  |  |  |  |
| GO:CC | GO:0005622 | intracellular | 2.15E-10 | GO:CC | GO:0031012 | extracellular matrix | 2.20E-13 |
| GO:CC | GO:0005737 | cytoplasm | 9.56E-10 | GO:CC | GO:0062023 | collagen-containing extracellular matrix | 6.96E-13 |
| GO:CC | GO:0012505 | endomembrane system | 4.41E-08 | GO:CC | GO:0005576 | extracellular region | 2.43E-10 |
| GO:CC | GO:0043229 | intracellular organelle | 6.67E-07 | GO:CC | GO:0005615 | extracellular space | 6.24E-10 |
| GO:CC | GO:0031410 | cytoplasmic vesicle | 4.46E-06 | GO:CC | GO:0005604 | basement membrane | 9.31E-10 |
| GO:CC | GO:0097708 | intracellular vesicle | 4.46E-06 | GO:CC | GO:0071944 | cell periphery | 1.83E-09 |
| GO:CC | GO:0043226 | organelle | 4.46E-06 | GO:CC | GO:0005886 | plasma membrane | 4.45E-09 |
| GO:CC | GO:0031982 | vesicle | 6.12E-06 | GO:CC | GO:0009986 | cell surface | 1.00E-08 |
| GO:CC | GO:0110165 | cellular anatomical entity | 6.99E-06 | GO:CC | GO:0110165 | cellular anatomical entity | 3.50E-06 |
| GO:CC | GO:0005829 | cytosol | 1.57E-05 | GO:CC | GO:0005587 | collagen type IV trimer | 1.68E-05 |
| GO:CC | GO:0048471 | perinuclear region of cytoplasm | 0.000242631 | GO:CC | GO:0098651 | basement membrane collagen trimer | 1.68E-05 |
| **Molecular Function** |  |  |  |  |  |  |  |
| GO:MF | GO:0005515 | protein binding | 1.08E-10 | GO:MF | GO:0005515 | protein binding | 6.12E-10 |
| GO:MF | GO:0019899 | enzyme binding | 1.69E-05 | GO:MF | GO:0005488 | binding | 3.83E-09 |
| GO:MF | GO:0008092 | cytoskeletal protein binding | 0.000331085 | GO:MF | GO:0005201 | extracellular matrix structural constituent | 1.62E-08 |
| GO:MF | GO:0005488 | binding | 0.000363156 | GO:MF | GO:0008201 | heparin binding | 1.58E-07 |
| GO:MF | GO:0071813 | lipoprotein particle binding | 0.026237583 | GO:MF | GO:0060089 | molecular transducer activity | 1.90E-07 |
| GO:MF | GO:0071814 | protein-lipid complex binding | 0.026237583 | GO:MF | GO:0005102 | signaling receptor binding | 3.41E-07 |
| GO:MF | GO:0001067 | regulatory region nuc. acid binding | 0.040927289 | GO:MF | GO:0005198 | structural molecule activity | 3.96E-07 |
| GO:MF | GO:0043168 | anion binding | 0.040927289 | GO:MF | GO:0005539 | glycosaminoglycan binding | 6.98E-07 |
| GO:MF | GO:0016740 | transferase activity | 0.040927289 | GO:MF | GO:0038023 | signaling receptor activity | 6.98E-07 |
| GO:MF | GO:0019900 | kinase binding | 0.040927289 | GO:MF | GO:0004888 | transmembr. signaling receptor activity | 1.04E-06 |
|  |  |  |  | GO:MF | GO:0035326 | cis-regulatory region binding | 2.38E-06 |
| **KEGG Pathway** |  |  |  |  |  |  |  |
| KEGG | KEGG:04810 | Regulation of actin cytoskeleton | 0.003951158 | KEGG | KEGG:04512 | ECM-receptor interaction | 1.60E-08 |
| KEGG | KEGG:04510 | Focal adhesion | 0.004660295 | KEGG | KEGG:04151 | PI3K-Akt signaling pathway | 2.72E-06 |
| KEGG | KEGG:04066 | HIF-1 signaling pathway | 0.004730602 | KEGG | KEGG:05165 | Human papillomavirus infection | 1.00E-05 |
| KEGG | KEGG:05205 | Proteoglycans in cancer | 0.004730602 | KEGG | KEGG:04360 | Axon guidance | 1.08E-05 |

|  |  |  |  |  |  |  |  |  |
| --- | --- | --- | --- | --- | --- | --- | --- | --- |
| KEGG | KEGG:01212 | Fatty acid metabolism | 0.006364804 |  | KEGG | KEGG:05200 | Pathways in cancer | 1.08E-05 |
| KEGG | KEGG:00051 | Fructose and mannose metabolism | 0.008062545 |  | KEGG | KEGG:04390 | Hippo signaling pathway | 3.33E-05 |
| KEGG | KEGG:05165 | Human papillomavirus infection | 0.00836325 |  | KEGG | KEGG:05146 | Amoebiasis | 0.000184329 |
| KEGG | KEGG:04140 | Autophagy - animal | 0.00836325 |  | KEGG | KEGG:05206 | MicroRNAs in cancer | 0.000205893 |
| KEGG | KEGG:05223 | Non-small cell lung cancer | 0.00836325 |  | KEGG | KEGG:05205 | Proteoglycans in cancer | 0.000273917 |
| KEGG | KEGG:04934 | Cushing syndrome | 0.008904189 |  | KEGG | KEGG:04550 | Signaling path. pluripotency of stem cells | 0.000328595 |
| KEGG | KEGG:05206 | MicroRNAs in cancer | 0.010714853 |  | KEGG | KEGG:04350 | TGF-beta signaling pathway | 0.000402713 |
| KEGG | KEGG:05214 | Glioma | 0.012951127 |  | KEGG | KEGG:01100 | Metabolic pathways | 0.000415169 |
| KEGG | KEGG:05163 | Human cytomegalovirus infection | 0.013108615 |  | KEGG | KEGG:04510 | Focal adhesion | 0.000454709 |
| KEGG | KEGG:04152 | AMPK signaling pathway | 0.01346403 |  | KEGG | KEGG:01230 | Biosynthesis of amino acids | 0.000852464 |
| KEGG | KEGG:05215 | Prostate cancer | 0.01346403 |  | KEGG | KEGG:05217 | Basal cell carcinoma | 0.001326581 |

**Table 2.**
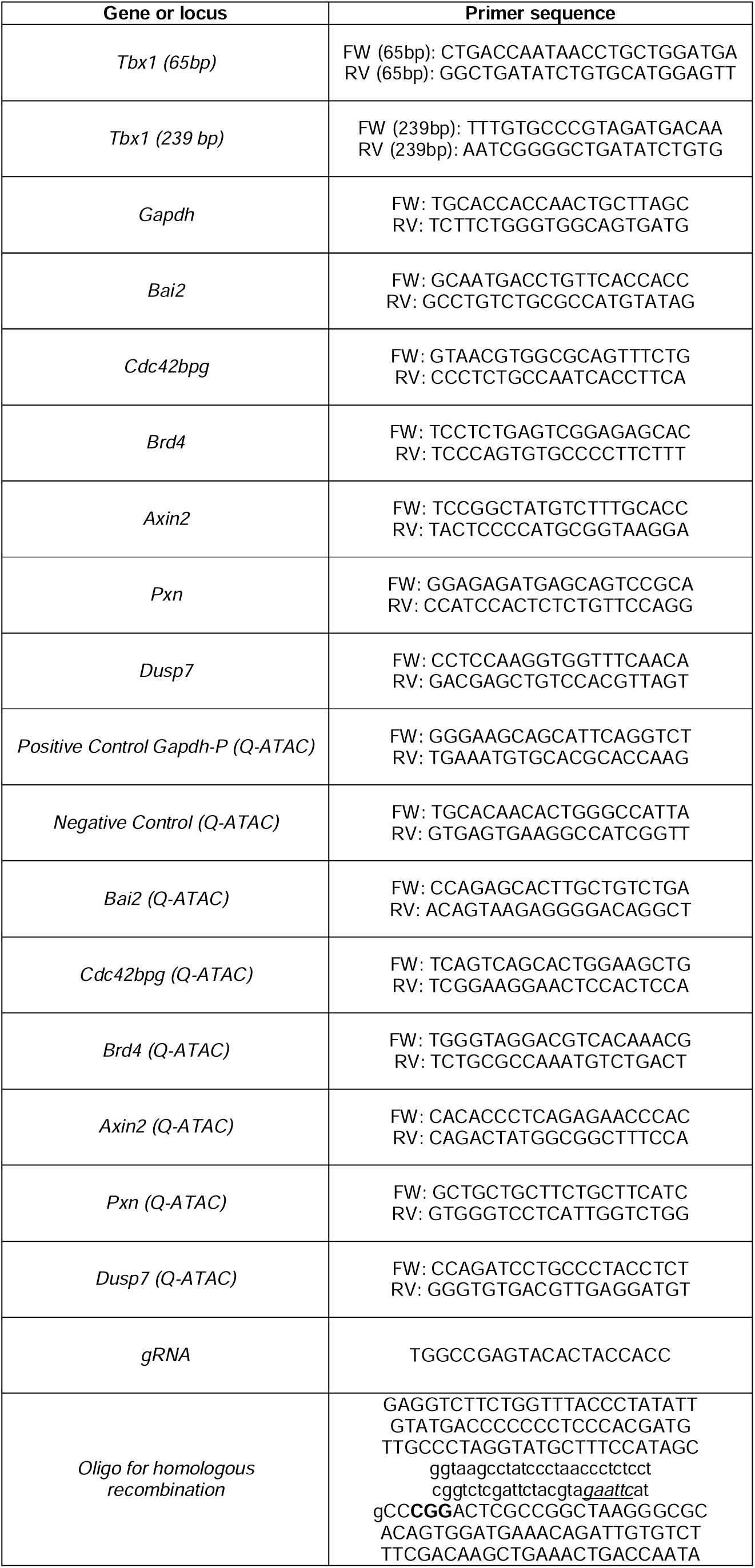
(primer sequences)

### ATAC-seq assay

Differentiated P19Cl6 cells and mESCs were collected and then washed two times in PBS, harvested, counted using a haemacytometer chamber and pelleted. 50.000 cells/sample for P19Cl6 and 15.000 cells/sample for mESC were treated with Tagment DNA Buffer 2x reaction buffer with Tagment DNA Enzyme (Illumina) according to the manufacturer’s protocol. After washes in PBS, cells were suspended in 50 μL of cold lysis buffer (10mM Tris-HCl, pH 7.4, 10mM NaCl, 3mM MgCl2, 0.1% IGEPAL CA-630) and immediately spun down at 500 x g for 10 min at 4 °C. Fresh nuclei were treated with Transposition mix and Purification (Illumina #FC121-130), the nuclei were incubated at 37 °C in Transposition Reaction Mix (25 μL reaction buffer, 2.5 μL Transposase, 22.5 μL Nuclease free water), purified using Qiagen MinElute PCR Purification Kit (Cat No./ID: 28006) and eluted in 10 μL of nuclease free water. Sequencing libraries were prepared from linearly amplified tagmented DNA. Fragmentation size was evaluated using the Agilent 4200 TapeStation. We sequenced two biological replicates for each experimental point. Sequencing was performed with an Illumina NextSeq500 machine, in paired-end, 60bp reads.

### Quantitative PCR ATAC (qATAC)

P19Cl6 cells were plated at a density of 5×10^5^ cells per well on a 35-mm tissue culture dish containing 25 pmol of a pool of Silencer Select Pre-Designed *Tbx1* SiRNA (Life Technology), non-targeted control (Life Technology) and 7.5 μl of RNAiMAX Reagent (Life Technology) diluted in 500 μl of Opti-MEM Medium (TermoFischer #31985062). We collected samples at 3 time points (see P19Cl6 paragraph above and scheme on Fig. 4A). For each time point we assayed two biological replicates. Cells were collected, washed, trypsinized and counted. Chromatin from 5×10^4^ cells was then tagmented, purified and used for quantitative PCR evaluation. To this end, we have used loci bound by TBX1 (Fulcoli et al., 2016) and located in open chromatin. For real-time PCR, we used biological duplicates for each time point and each duplicate was divided into two technical replicates. Two different controls were used: *Gapdh* promoter (positive control) representing open chromatin, and a desert island locus (negative control), which does not contain any genes in a range of about 80 kb. Quantification was performed using 2^-ΔCt calculation relative to *Gapdh* promoter (positive control). The data are expressed as the average of two biological replicates and the standard deviation. Primer sequences are reported on Tab. 2.

**Figure 2.**
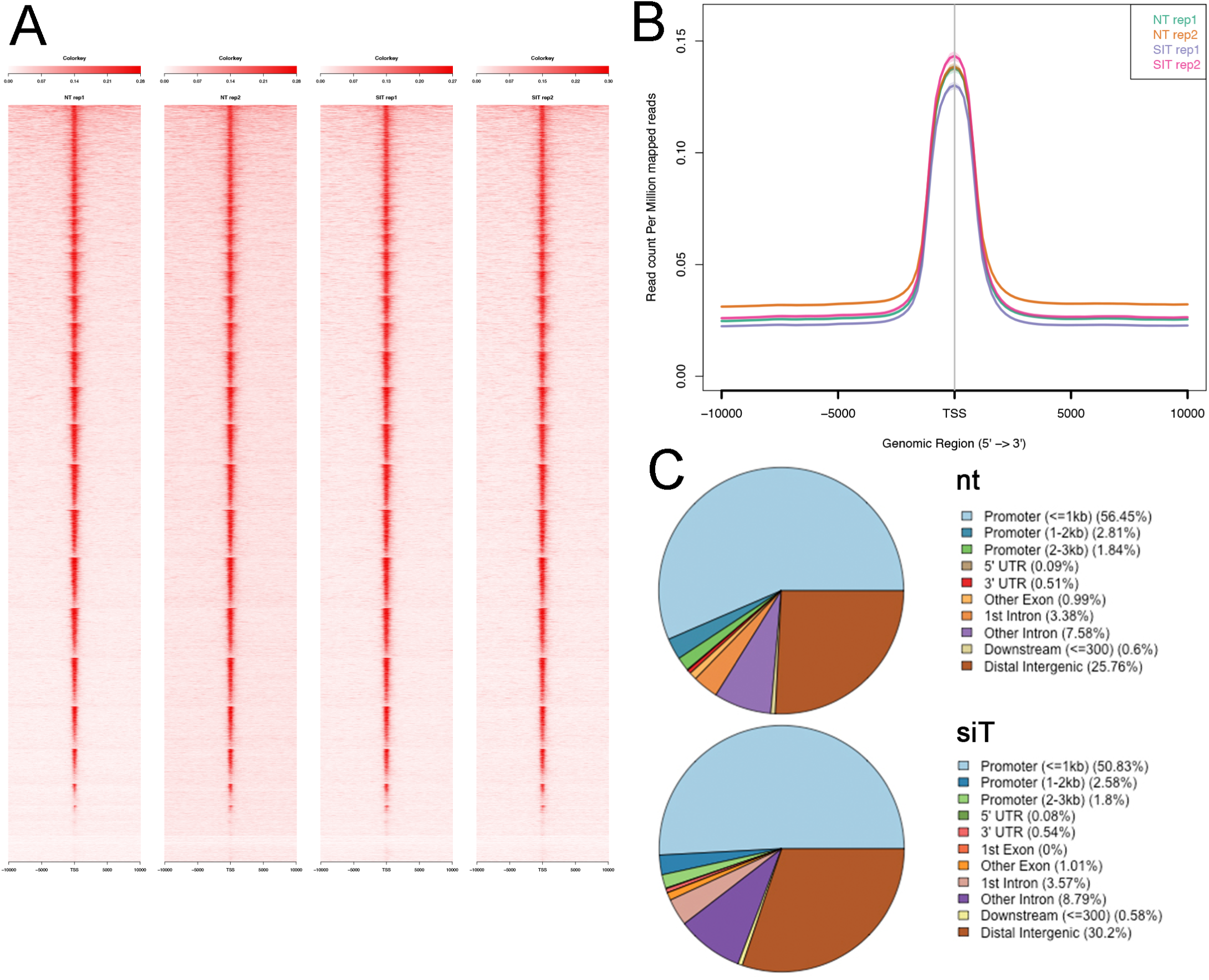
ATAC-seq data using P19Cl6 cells differentiated at Day 1. A) Heat maps of signal distribution around the transcription start site (TSS) +/- 10000bp of 14476 expressed genes in two control (NT, non-targeted siRNA) and two *Tbx1*^*KD*^ (SIT) replicates. B) Average profiles of enrichment at the TSS in control and *Tbx1*^*KD*^ cells. Note the similar distribution of all 4 samples. C) Pie charts illustrate the distribution of ATAC-seq consensus peaks relative to gene features. The distributions are very similar in controls (nt, number of consensus peaks= 23759) and *Tbx1*^*KD*^ cells (siT, number of consensus peaks= 26847).

**Figure 3.**
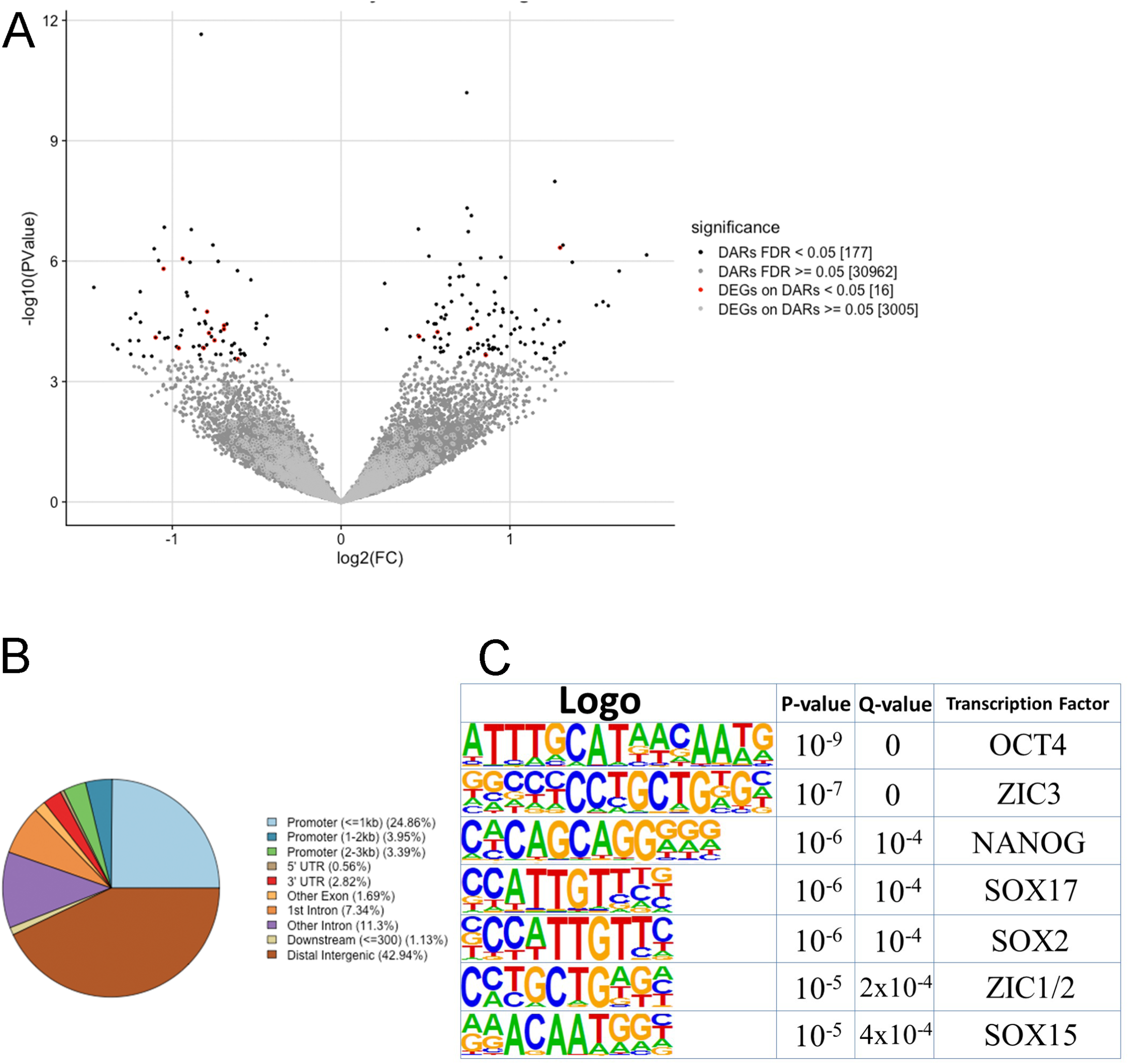
Analysis of differentially accessible regions (DARs) in *Tbx1*^*KD*^ vs. control P19Cl6 cells. A) Volcano plot of all peaks. Regions with significantly different accessibility are indicated as black dots. Red dots indicate DARs associated with differentially expressed genes (DEGs). B) Pie chart showing the distribution of the 177 DARs relative to gene features. Note the reduced representation of promoter regions and a relatively higher representation of distal intergenic regions compared to the general populations of peaks shown in Fig. 2C. C) Logos of the most significantly enriched motifs detected in the 177 DARs.

**Figure 4.**
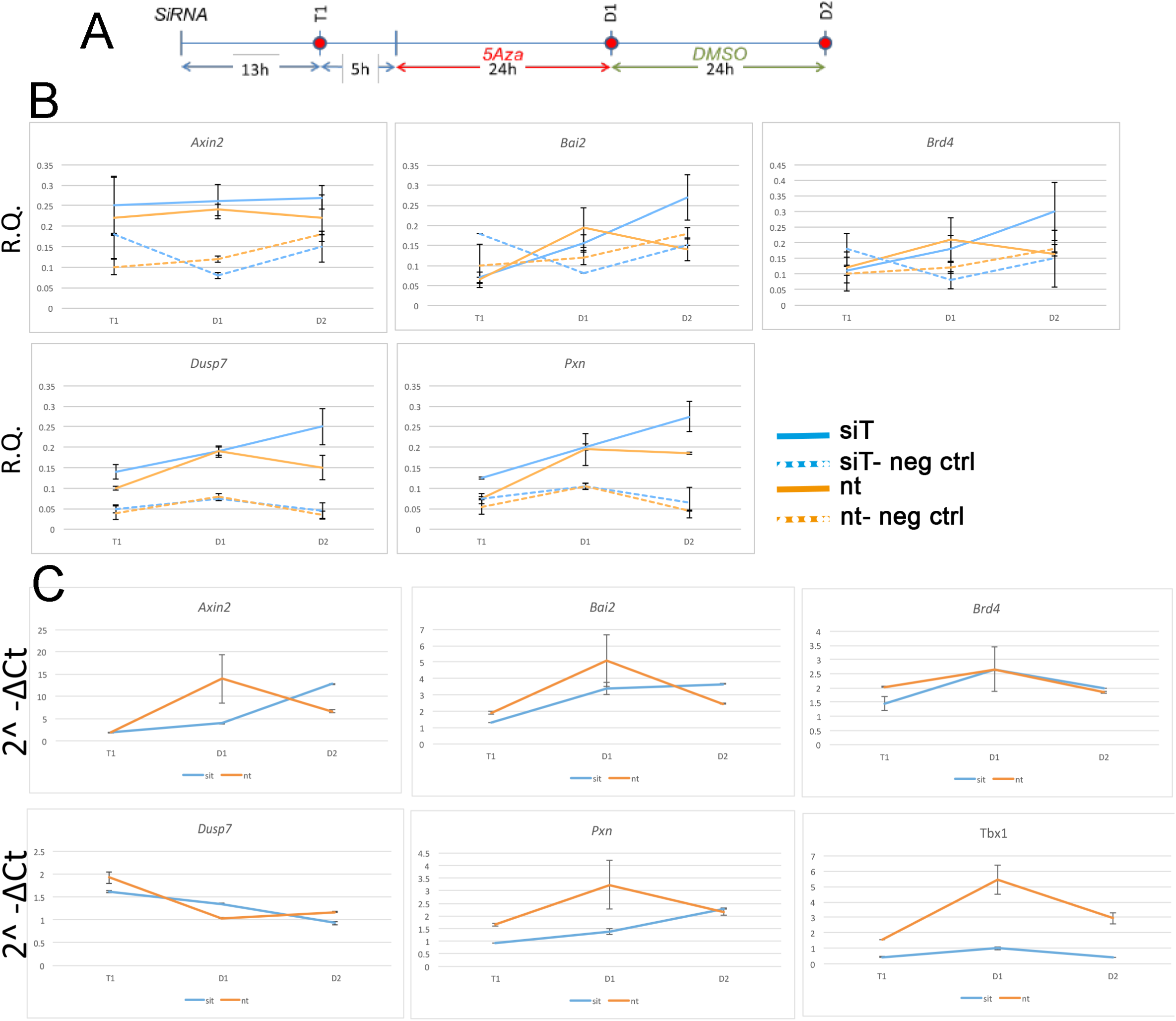
Changes in chromatin accessibility during differentiation in P19Cl6 cells. A) Experimental scheme illustrating the three time points tested, T1 (13hrs after transfection of siRNA), D1 (24 hrs after 5Aza addition to the media), and D2 (24 hrs after addition of DMSO to the media). B) Quantitative ATAC assays of previously identified TBX1 binding sites associated with the genes indicated. All sites were found to be accessible by ATAC-seq. In all cases, accessibility tends to increase at D2. The negative control locus is located in a gene desert region (see Tab. 2 for primers sequences). Values are the average of two biological replicates +/- standard deviation. C) Gene expression analysis by quantitative real time PCR of the same genes. Values are the average of two biological replicates +/- standard error of the mean.

RNA was also extracted from each sample and the expression of genes associated with the above loci was evaluated using real-time reverse transcription PCR (qRT-PCR). Quantitation was performed using relative quantification (RQ) and calculated with the standard error of the mean of two biological replicates.

### RNA-seq

mESCs in dishes were washed with cold PBS to which 1 mL of Trizol was added. Lysates were harvested and vortexed in order to promote the lysis of cells. 200 μL of chloroform was added to 1 mL of lyste in order to separate the phases. The mixture was centrifuged at 12000g for 15 min. The upper phase was removed and transferred into a new tube containing 500 μL of isopropanol and the solution was incubated for 20 min. at room temperature. After 20 min. the solution was centrifuged for 10 min. at 12000g. Pelleted RNA was washed twice with Ethanol 80% and centrifuged for 5 min. at 7500g. Pellet were resuspended, and the concentration was estimated using a Nanodrop. Libraries were prepared according to the Illumina strand specific RNA-seq protocol. Libraries were sequenced on the Illumina platform NextSeq 500, in paired end, 75bp reads.

### RNA-seq and ATAC-seq data analysis

Expressed and differentially expressed (DE) genes related to the analysis of the RNA-seq samples in the P19Cl6 cellular model were retrieved from published datasets (Fulcoli et al., 2016). Mouse ESCs RNA-seq raw sequences were first evaluated for quality using FastQC (https://www.bioinformatics.babraham.ac.uk/projects/fastqc/), then mapped to the mouse genome (mm9) using TopHat2 2.0.14 (Kim et al 2013) with -*r 170 --mate-std-dev 50 --transcriptome-index transcriptome --library-type fr-secondstrand -N 3 --read-edit-dist 5* and all other parameters as default. The transcriptome was built using the Mus_musculus.NCBIM37.67.gtf annotation downloaded from the ensemble database (http://www.ensemble.org). Only uniquely mappable sequences were retained for further analysis. For each sample, the gene expression was quantified in terms of raw counts using HTseq 0.7.1 (Anders et al., 2015) with *-m intersection-nonempty -s reverse* for all annotated genes. The next analysis was carried out using RNASeqGUI 1.2.1 (Russo et al., 2016), where the expressed genes were first selected using the proportion test, then the raw counts were normalized using the upper quartile method. Finally, the DESeq2 module was used to identify differentially expressed (DE) genes. Genes with adjusted p-values <0.05 were considered DE. Pathway analysis was carried out for both cellular models using gprofiler2 (Raudvere et al., 2019), with the expressed genes as background and a threshold of 0.05 for the FDR value.

For ATAC-seq analysis, FastQC quality check showed 10-20% contamination of Nextera Transposase Sequence primers (Turner, 2014) in the range 33 to 47 bp. We removed these sequences using cutadapt (Martin, 2011) with the following option -a CTGTCTCTTATACACATCTCCGAGCCCACGAGAC -A CTGTCTCTTATACACATCTGACGCTGCCGACGA. Sequences were then aligned to the mouse genome (mm9) using Bowtie2 2.3.4.3 (Langmead and Salzberg, 2012) with the default parameters. Only uniquely mappable reads were retained. A customized R script was used to remove reads with mates mapping to different chromosomes, or with discordant pairs orientation, or with a mate-pair distance >2 kb, or PCR duplicates (defined as when both mates are aligned to the same genomic coordinate). Reads mapping to the mitochondrial genome were also removed. Coverage heat-maps and average enrichment profiles (TSS +/- 10Kb) in each experimental condition were obtained using ngs.plot (Shen et al., 2014) and evaluated on the expressed genes of the cellular models.

ATAC peaks were identified using MACS2 2.1.2.1 (Feng et al., 2012) with the option *-- nomodel --shif100 --extsize 200* that are the suggested parameters to handle the Tn5 transposase cut site. In particular, peak calling was performed independently on each ATAC-seq sample of the P19Cl6 cellular model. Then, for each condition a consensus list of enriched regions was obtained using the intersectBed function from the BedTools 2.29 (Quinlan and Hall, 2010), with the default minimum overlap and retaining only the peak regions common to both replicates. In contrast, for the mouse ES cellular model, due to the lower coverage, the two replicates were first merged into a single signal, after which MACS2 was applied to the pooled samples for each experimental condition.

For the P19Cl6 ATAC cellular model, unless otherwise specified, differentially enriched regions (DARs) were obtained using DEScan2 1.6.0 (Righelli et al., 2018) by loading all of the MACS2 peaks, and performing the peak consensus with the finalRegions function (zThreshold=1, minCarriers=2 parameters) and using edgeR with estimateDisp, glmQLFit (robust=TRUE parameter), glmQLFTest, in this order and with the defaults parameters. For the pooled mouse ES samples, the function sicer_df (Zang et al., 2009) was used to identify the DARs, setting 200, 400, 0.00001, 0.0001 as parameters for window size (bp), gap size (bp), and FDR_vs_Input, FDR, respectively.

Peaks, consensus peaks, and DARs were annotated with genes using ChIPseeker 1.22 (Yu et al., 2015) by associating to each peak/region the nearest gene, setting the TSS region [-3000, 3000] and downloading the annotation “may2012.archive.ensembl.org” from Biomart (using the makeTxDbFromBiomart function) and the org.Mm.eg.db database. Finally, the annotated genes were intersected with the DE genes.

For both cellular models, transcription factor binding motifs were obtained using the findMotifsGenome program of the HOMER suite (http://homer.ucsd.edu/homer/).

Volcano plots, pie-charts, and other data reshaping were performed using standard R-scripts. Overlaps among different regions were identified using the intersectBed function from the BedTools 2.29, with default minimum overlap and reporting each original entry once.

## RESULTS

### Cell differentiation models

P19Cl6 cells were subjected to a differentiation protocol that has been previously described (Mueller et al., 2010) (Fig. 1A). Cells were transfected with scrambled or *Tbx1*-targeted siRNAs, harvested at day 1 of differentiation and processed for ATAC-seq in 2 biological replicates.

Mouse ES cells were targeted using CRISPR-Cas9 in order to generate a homozygous *Tbx1* loss of function mutation by inserting multiple stop codons and polyA signals into exon 5 by homologous recombination. We obtained 2 correctly targeted homozygous mutant clones and selected one of them (5H) for further experiments because it did not express any *Tbx1* mRNA (Fig. 1D-E). Clone 5H and the parental cell line (WT) were subjected to a differentiation protocol (Kattman et al., 2011) according to the scheme shown in Fig. 1A. WT cells expressed *Tbx1* at day 4, while no expression was detected in *Tbx1*^-/-^ cells (Fig. 1F).

To enrich for *Tbx1*-expressing cells, we subdivided the population of mES cells by FACS using the standard markers PDGFRA and KDR (a.k.a. VEGFR2) at day 4 of differentiation (Fig. 1B). We extracted RNA from sorted populations, KDR+;PDGFRa-, KDR+;PDGFRa+, KDR-;PDGFRa+, and from the total, unsorted population. qRT-PCR showed that by far the highest expression of *Tbx1* was in the KDR-;PDGFRa+ population (Fig. 1C). The same fractionation was performed on *Tbx1*^-/-^ 5H cells and we found that the quantitative distribution of cells in the fractions was similar (Fig. S1, S2). Therefore, ATAC-seq and RNA-seq assays were performed using KDR-;PDGFRa+ day 4 cells from the WT and *Tbx1*^-/-^ lines, in 2 biological replicates.

### Chromatin accessibility assay

#### A) P19Cl6 cells

Control (scrambled siRNA treated) and *Tbx1* depleted (*Tbx1*^*KD*^) cells exhibited a similar distribution and intensity of ATAC-seq signal, which was mostly localized to the promoter region of genes, as expected (Fig. 2A-C).

We next compared ATAC-seq data with TBX1 ChIP-seq data previously reported for this cell line, and under the same differentiation conditions (Fulcoli et al., 2016). Surprisingly, we found that only 80 ATAC peaks (out of a total of 23759 non-targeted siRNA peaks) overlapped with 72 TBX1 ChIP-seq peaks (3% of the 2388 TBX1 peaks), indicating that most TBX1 binding sites are located in closed chromatin, i.e. ATAC-negative.

We next compared chromatin accessibility profiles in control and *Tbx1*^*KD*^ cells in order to identify differentially accessible regions (DARs) between the two conditions. We found a total of 177 DARs, of which, 72 (41%) had reduced accessibility and 105 (59%) had increased accessibility following *Tbx1* knockdown (Fig. 3A). The 177 DARs were annotated with 165 distinct genes, according to the gene symbol (Supplementary Tab. 1). Comparison of this gene list with previously identified differentially expressed genes (DEGs) (Fulcoli et al., 2016) revealed that only 16 (9%) were differentially expressed (Fig. 3A, list in Supplementary Tab. 1); thus, in most cases, chromatin changes identified in our dataset were not associated with transcriptional changes measured by RNA-seq. We examined the distribution of DARs relative to gene features and found that compared to the distribution of all peaks in the WT population (Fig. 2C), there was a relatively low presence in the promoter regions (32.2% vs. 61.1%) and a relatively higher representation in distal intergenic regions (42.9% vs. 25.8%).

Analysis of DAR sequences identified a set of known motifs of transcription factors with homeodomains (OCT4, NANOG), high mobility group domains (SOX2, SOX17, SOX15), and zinc finger domains (ZIC1, ZIC2, ZIC3) (Fig. 3C). Interestingly, several of these proteins are pluripotency factors, but we did not detect any enrichment of T-BOX binding motifs. This is consistent with the finding that only one of all of the DARs identified here overlapped with TBX1 ChIP-seq peaks (indicated in Supplementary Tab. 1).

The finding that almost no TBX1 ChIP-seq peaks changed accessibility after loss of TBX1 was very surprising to us. In order to test whether accessibility changes might follow *Tbx1* KD at a later time than the one tested here, we performed a time-course experiment on 5 ChIP-seq peaks located in open chromatin and associated with 5 target genes (Fulcoli et al., 2016). The experimental scheme is shown in Fig. 4A. At each time point, we carried out quantitative ATAC (qATAC) in control and *Tbx1*^*KD*^ cells. At each point, we also measured the expression of the target genes. The results (Fig. 4B-C), which were normalized for the accessibility at the GAPDH promoter, confirmed that at D1 (the time tested by ATAC-seq) there was no change in chromatin accessibility. However, at D2, 4 out of 5 loci showed increased accessibility in the *Tbx1*^*KD*^ condition. In all cases, gene expression changed at earlier time points (T1 or D1, Fig. 4C), suggesting that differential expression, for these genes, preceded chromatin changes. These results suggest that the effects of *Tbx1* KD on chromatin accessibility are most likely to be indirect.

#### B) Mouse ES cells

We performed ATAC-seq assays on two biological replicates of differentiating WT and *Tbx1*^-/-^ mES cells selected by FACS (PDGFRA+;KDR-). The distribution of ATAC-seq peaks was similar between control and mutant cells (Fig. 5A-C). We found a total of 138 DARs, of which 26 (19%) decreased accessibility, and 112 (81%) increased accessibility in *Tbx1*^-/-^ cells. Their distribution, compared to WT peaks distribution (Fig. 5C) showed a relatively lower representation in the promoter regions (43.5% vs. 75.2%) and relatively higher representation in the distal intergenic regions (31.2% vs. 15%), similarly to what we found in P19Cl6 DARs. The 138 peaks were annotated with 133 distinct genes, according to the gene symbol (Supplementary Tab. 3), of which 12 (8.7%) were also differentially expressed (Supplementary Tab. 3). Sequence analysis of DARs revealed an enrichment of T-BOX binding motifs (Fig. 5D), suggesting that TBX1 or other T-BOX proteins might occupy some of these sites. The two T-BOX motifs shown in Fig. 5D are almost identical, and one or the other was found in 47 DARs (34% of all DARs) (Supplementary Tab. 3, T-BOX column). Interestingly, 43 out of 47 (91%) were more accessible in mutant cells, suggesting that in these cells, TBX1 may work to maintain the chromatin close at selected loci, consistent with results obtained in the time course experiment with P19Cl6 cells.

**Figure 5.**
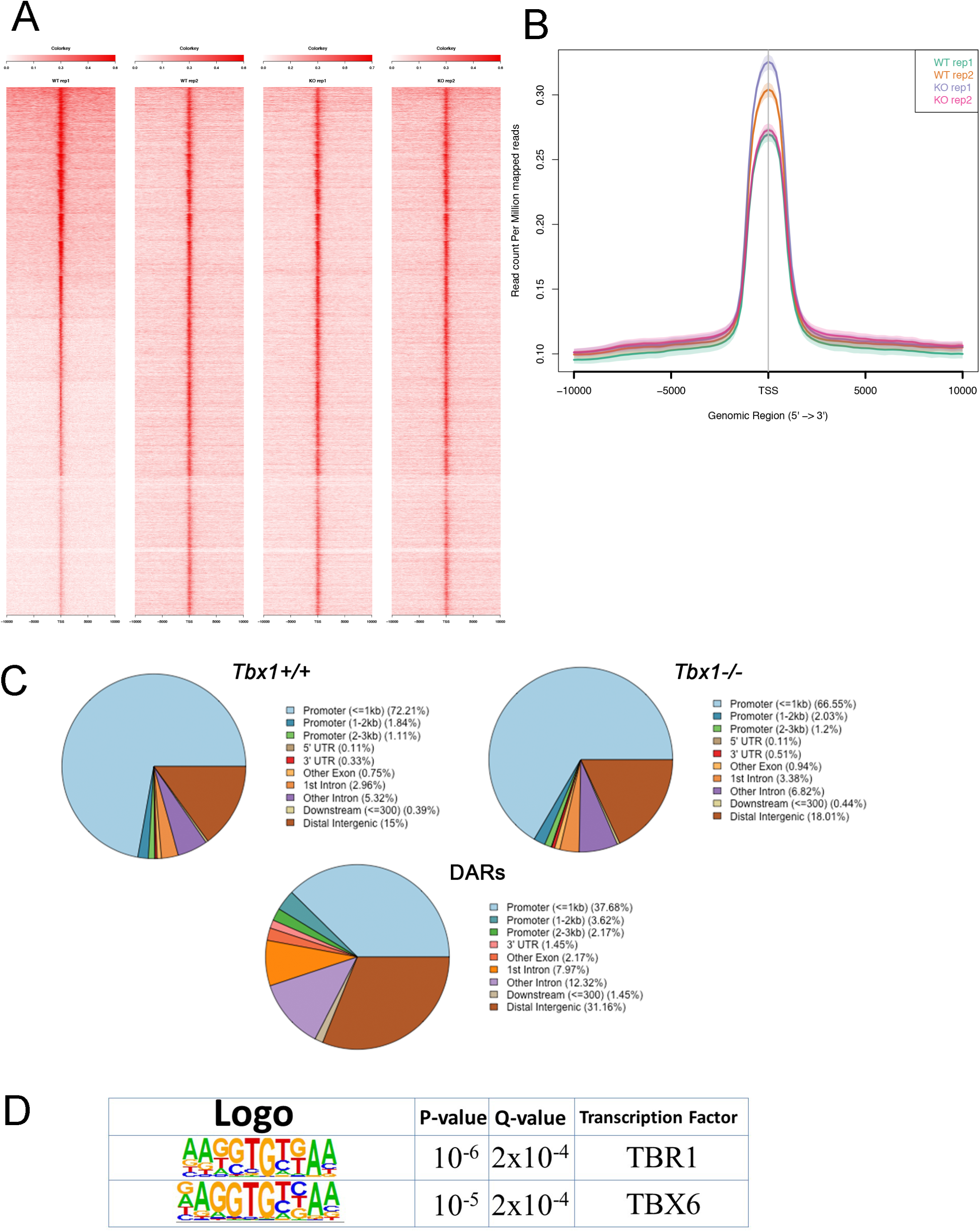
ATAC-seq of PDGFRA+;KDR-differentiating mESCs at Day 4 of differentiation. A) Heat maps of signal distribution around the transcription start site (TSS) +/- 10000bp of the 13075 expressed genes in two *Tbx1*^+/+^ (WT) and two *Tbx1*^*-/-*^ (KO) biological replicates. B) Average profiles of enrichment at the TSS of the 13075 expressed genes in WT and KO cells. C) Pie charts of the distribution of ATAC-seq peaks in WT (top left, number of peaks=11362), KO (top right, number of peaks=13622), and DARs (bottom, n=138) relative to gene features. Note that as for P19Cl6, the DARs are relatively less enriched in the promoter region, and more enriched in the distal intergenic regions. D) Logos of the most enriched known motifs in the 138 DARs. The two consensus sequences are almost identical and reproduce a typical T-BOX binding motif.

### Transcriptional profiling

Transcriptional changes in response to *Tbx1* knockdown have been reported for P19Cl6 cells (Fulcoli, 2016). Here, we performed RNA-seq analysis on WT and *Tbx1*^-/-^ cells that derive from the same differentiation experiments (2 biological replicates) as the ATAC-seq experiments described above. Results revealed 642 genes to be differentially expressed; 412 down-regulated, 230 up regulated in *Tbx1*^-/-^ cell line compared to the parental WT cell line in two biological replicates (Supplementary Tab. 2). A list of all genes expressed in these cells is shown in Supplementary Tab. 4. Next, we carried out functional profiling/gene ontology analyses with g:Profiler2 using DEGs from mES cells and from P19Cl6 cells (Fulcoli et al., 2016) using identical criteria. Results are shown side-by-side on Tab. 1. Results were very similar in the two models for the gene ontology category “biological process” because in both cases, DEGs were enriched in developmental genes. However, we noted substantial differences in the category of “cellular component” where mESC showed strong enrichment of genes related to the extracellular matrix (ECM), while in contrast, P19Cl6 cells showed enrichment for genes related to intracellular components. KEGG pathway analysis showed again a strong enrichment of ECM-related pathways but with some limited overlap with the P19Cl6 results, as both models showed focal adhesion to be among the enriched pathways. The enrichment of the KEGG pathways “ECM-receptor interaction” and “basal cell carcinoma” categories in the mESC model is also consistent with recent findings in the mouse (Alfano et al., 2019; Caprio et al., 2020).

## DISCUSSION

Data analysis of two cell culture models provided a snapshot of the chromatin accessibility, as measured by ATAC-seq, with and without TBX1 function, or dosage reduction. The use of cell culture systems has limitations because they do not mimic complex developmental processes, but they also have some advantages because they are relatively homogeneous compared to whole-organ or whole-embryo material. This is particularly true for our mESC model in which we used a specific subpopulation at a specific differentiation point. In both models, loss of TBX1 led to significant transcriptional changes, as measured by RNA-seq, indicating that both models respond robustly to loss or reduced dosage of TBX1.

In differentiating P19Cl6 cells, the intersection of ATAC signals with a map of TBX1 binding sites, which was previously published for the same cell line and under the same experimental conditions, revealed that almost all of the binding sites were located in ATAC-negative regions. Thus, TBX1 binding does not require accessible chromatin, at least by ATAC assay. Unfortunately, we were not able to confirm this finding in differentiating mES cells because in our hands, currently available batches of commercial anti-TBX1 antibodies failed to perform in ChIP experiments. In future experiments it will be of interest to establish whether TBX1 can function as a pioneer factor. In a recent paper, it was shown that TBX20, a T-BOX transcription factor that belongs to the same sub-family as TBX1, mostly binds (2/3 of the cases) in closed chromatin regions in endocardial cells (Boogerd et al., 2016). Thus, T-BOX proteins may not need ATAC+ regions to bind chromatin. It is also possible that TBX1, at early stages of differentiation, contributes to maintaining the chromatin closed at selected loci. Indeed, in mESC experiments almost all the DARs with a T-BOX binding motif showed increased accessibility in *Tbx1*^-/-^ cells.

Gene ontology (GO) analysis of DEGs in response to loss of TBX1 in the two models revealed broad differences, but also some similarities. In both cases, DEGs were enriched in genes involved in developmental processes, as expected; in both cases, the focal adhesion KEGG pathway was significantly enriched. The latter has been validated recently in different cultured cells and in mouse mutants (Alfano et al., 2019). However, in general, GO enrichment was more dispersed in P19Cl6 cells compared to differentiated mES cells, where there was higher enrichment of specific GO terms, perhaps reflecting a more differentiated state and/or a more homogeneous cell population. Particularly evident was the presence of ECM-related genes in the mESC-derived cells.

We selected PDGFRA+;KDR-cells for our studies on the basis of *Tbx1* gene expression; data in the literature suggest that mESC-derived PDGFRA+;KDR-cells have a marker profile that is similar to paraxial mesoderm (Craft et al., 2013; Sakurai et al., 2006; Tanaka et al., 2009), which includes head mesenchyme, a tissue that expresses high levels of *Tbx1*. The mesenchymal nature of the PDGFRA+;KDR-cell population is also consistent with high expression of genes encoding collagens and vimentin. *Tbx1*-expressing mesenchymal cells contribute to various tissues of the neck and face, including some muscle, bones, and connective tissue (Adachi et al., 2020).

Despite a significant transcriptional response to loss of TBX1, we detected a very modest chromatin response in both models. We have previously proposed that TBX1 is a priming factor that regulates deposition of H3K4me1 in H3K27Ac-negative regions, presumed to be inactive enhancers (Fulcoli et al., 2016). This hypothesis is consistent with our finding of closed chromatin at most TBX1-binding sites. However, H3K4me1 triggers a number of mechanisms that eventually lead to the opening of the chromatin (reviewed in (Calo and Wysocka, 2013)). Thus, we would have expected a more extensive chromatin remodeling than the one that we observed. However, H3K4me1 deposition may also lead to long range chromatin changes, which were not tested in our experiments (Yan et al., 2018). In any case, it is possible that the putative priming activity of TBX1 leads to chromatin changes that are downstream of the differentiation time window tested here. A recent study tested the effect of FOXA2 loss of function on chromatin remodeling during ES-based endoderm differentiation into pancreatic cells (Lee et al., 2019). FOXA2 is a pioneer transcription factor that, like TBX1, regulates H3K4me1 deposition in early phases of differentiation (to definitive endoderm), but its loss did not cause significant ATAC-seq changes at this stage. It was only at later stages of differentiation, when H3K27 acetylation occurred, that ATAC-seq changes became significant (Lee et al., 2019). Our qATAC time course results, though limited to a small number of loci, is consistent with the hypothesis that chromatin remodeling may occur at later stages of differentiation. The development of optimized protocols that will allow the monitoring of *Tbx1*-expressing cells throughout their differentiation will help to address this hypothesis in the future.

## ACKNOWLEDGMENTS

This work was funded in part by grants from the Italian Ministry of Research PRIN 20179J2P9J (to AB, EI, GL, and CA), the Campania SATIN Project (to AB), and the Fondation Leducq grant 15CVD01 (to AB and EI). We wish to thank the FACS core of the Institute of Genetics and Biophysics for their assistance with cell sorting.

## Author Contributions

AC, IA, GL, RF, performed wet lab experiments. AC, MF, DR, CA, performed bioinformatics data analysis, EI, CA, AB provided funds, AB wrote the manuscript, EI, AC, IA, GL, CA contributed to writing and editing.

## SUPPLEMENTARY MATERIAL

**Supplementary Figure 1**

Analysis of surface markers PDGFRA and KDR expression in *Tbx1*^+/+^ and *Tbx1*^-/-^ mESC lines at day 4 of differentiation. A, D) *Tbx1*^+/+^ and *Tbx1*^-/-^ cells labeled by primary antibodies PDGFRA (APC-A) and KDR (PE-A). 4 fractions were identified: Q1-2 = PDGFRA+/KDR-; Q2-2 = PDGFRA+/KDR+; Q4-2= PDGFRA-/KDR+; Q3-2= PDGFRA-/KDR-; B-C-E-F) Negative controls (NC) are cells not incubated with primary antibodies.

**Supplementary Figure 2**

Plots showing FACS controls. A) *Tbx1*^+/+^ cells labeled by PDGFRA-APC antibody and isotype control-PE (left); and by isotype control-APC and KDR-PE antibodies (right); B) *Tbx1*^-/-^ cells labeled by PDGFRA-APC antibody; and isotype control-PE (left), by isotype control-APC and KDR-PE antibodies (right).

**Supplementary Table 1**

Differentially accessible regions in P19Cl6 cells with gene annotation, intersection with differentially expressed genes, and intersection with TBX1 ChIP-seq peaks.

**Supplementary Table 2**

Differentially expressed genes in differentiating mESC. Columns contain the Ensembl Identifier, the corresponding Gene Symbol, the log2 FoldChange of KO/WT and the Adjusted P-value (cut off =0.05).

**Supplementary Table 3**

Differentially accessible regions in differentiating mES cells with gene annotation, intersection with differentially expressed genes.

**Supplementary Table 4**

Genes expressed in PDGFRA+;KDR-mES cells at d4. Columns contain the Ensembl Identifier, the corresponding Gene Symbol and the normalized upper quartile gene expression counts for each sample.

## Notes

### Competing Interest Statement

The authors have declared no competing interest.

### Summary of Updates

The T-box transcription factor TBX1 has critical roles in the cardiopharyngeal lineage and the gene is haploinsufficient in DiGeorge syndrome, a typical developmental anomaly of the pharyngeal apparatus. Despite almost two decades of research, if and how TBX1 function triggers chromatin remodeling is not known. Here, we explored genome-wide gene expression and chromatin remodeling in two independent cellular models of Tbx1 loss of function, mouse embryonic carcinoma cells P19Cl6, and mouse embryonic stem cells (mESCs). The results of our study revealed that the loss or knockdown of TBX1 caused extensive transcriptional changes, some of which were cell type-specific, some were in common between the two models. However, unexpectedly we observed only limited chromatin changes in both systems. In P19Cl6 cells, differentially accessible regions (DARs) were not enriched in T-BOX binding motifs; in contrast, in mESCs, 34% (n=47) of all DARs included a T-BOX binding motif and almost all of them gained accessibility in Tbx1-/- cells. In conclusion, despite a clear transcriptional response of our cell models to loss of TBX1 in early cell differentiation, chromatin changes were relatively modest.

## REFERENCES

Adachi, N., Bilio, M., Baldini, A. and Kelly, R. G. (2020). Cardiopharyngeal mesoderm origins of musculoskeletal and connective tissues in the mammalian pharynx. Dev. Camb. Engl. 147,.

Alfano, D., Altomonte, A., Cortes, C., Bilio, M., Kelly, R. G. and Baldini, A. (2019). Tbx1 regulates extracellular matrix-cell interactions in the second heart field. Hum. Mol. Genet. 28, 2295–2308.

Anders, S., Pyl, P. T. and Huber, W. (2015). HTSeq--a Python framework to work with high-throughput sequencing data. Bioinforma. Oxf. Engl. 31, 166–169.

Baldini, A., Fulcoli, F. G. and Illingworth, E. (2017). Tbx1: Transcriptional and Developmental Functions. Curr. Top. Dev. Biol. 122, 223–243.

Boogerd Cj, Aneas I, Sakabe N, Dirschinger Rj, Cheng Qj, Zhou B, Chen J, Nobrega Ma and Evans Sm (2016). Probing Chromatin Landscape Reveals Roles of Endocardial TBX20 in Septation. J. Clin. Invest. 126,.

Buenrostro, J. D., Wu, B., Chang, H. Y. and Greenleaf, W. J. (2015). ATAC-seq: A Method for Assaying Chromatin Accessibility Genome-Wide. Curr. Protoc. Mol. Biol. 109, 21.29.1-21.29.9.

Calo, E. and Wysocka, J. (2013). Modification of enhancer chromatin: what, how, and why? Mol. Cell 49, 825–837.

Caprio, C., Varricchio, S., Bilio, M., Feo, F., Ferrentino, R., Russo, D., Staibano, S., Alfano, D., Missero, C., Ilardi, G., et al. (2020). TBX1 and Basal Cell Carcinoma: Expression and Interactions with Gli2 and Dvl2 Signaling. Int. J. Mol. Sci. 21,.

Castellanos, R., Xie, Q., Zheng, D., Cvekl, A. and Morrow, B. E. (2014). Mammalian TBX1 pREFerentially binds and regulates downstream targets via a tandem T-site repeat. PloS One 9, e95151.

Chen, L., Fulcoli, F. G., Ferrentino, R., Martucciello, S., Illingworth, E. A. and Baldini, A. (2012). Transcriptional control in cardiac progenitors: Tbx1 interacts with the BAF chromatin remodeling complex and regulates Wnt5a. PLoS Genet. 8, e1002571.

Craft, A. M., Ahmed, N., Rockel, J. S., Baht, G. S., Alman, B. A., Kandel, R. A., Grigoriadis, A. E. and Keller, G. M. (2013). Specification of chondrocytes and cartilage tissues from embryonic stem cells. Dev. Camb. Engl. 140, 2597–2610.

Feng, J., Liu, T., Qin, B., Zhang, Y. and Liu, X. S. (2012). Identifying ChIP-seq enrichment using MACS. Nat. Protoc. 7, 1728–1740.

Fulcoli, F. G., Franzese, M., Liu, X., Zhang, Z., Angelini, C. and Baldini, A. (2016). Rebalancing gene haploinsufficiency in vivo by targeting chromatin. Nat Commun 7, 11688.

Greulich, F., Rudat, C. and Kispert, A. (2011). Mechanisms of T-box Gene Function in the Developing Heart. Cardiovasc. Res. 91,.

Kattman, S. J., Witty, A. D., Gagliardi, M., Dubois, N. C., Niapour, M., Hotta, A., Ellis, J. and Keller, G. (2011). Stage-specific optimization of activin/nodal and BMP signaling promotes cardiac differentiation of mouse and human pluripotent stem cell lines. Cell Stem Cell 8, 228–240.

Langmead, B. and Salzberg, S. L. (2012). Fast gapped-read alignment with Bowtie 2. Nat. Methods 9, 357–359.

Lee, K., Cho, H., Rickert, R. W., Li, Q. V., Pulecio, J., Leslie, C. S. and Huangfu, D. (2019). FOXA2 Is Required for Enhancer Priming during Pancreatic Differentiation. Cell Rep. 28, 382-393.e7.

Martin, M. (2011). Cutadapt removes adapter sequences from high-throughput sequencing reads. EMBnet.journal 17, 10–12.

McDonald-McGinn, D. M., Sullivan, K. E., Marino, B., Philip, N., Swillen, A., Vorstman, J. A. S., Zackai, E. H., Emanuel, B. S., Vermeesch, J. R., Morrow, B. E., et al. (2015). 22q11.2 deletion syndrome. Nat. Rev. Dis. Primer 1, 15071.

Mueller, I., Kobayashi, R., Nakajima, T., Ishii, M. and Ogawa, K. (2010). Effective and steady differentiation of a clonal derivative of P19CL6 embryonal carcinoma cell line into beating cardiomyocytes. J Biomed Biotechnol 2010, 380561.

Okubo, T., Kawamura, A., Takahashi, J., Yagi, H., Morishima, M., Matsuoka, R. and Takada, S. (2011). Ripply3, a Tbx1 repressor, is required for development of the pharyngeal apparatus and its derivatives in mice. Dev. Camb. Engl. 138, 339–348.

Quinlan, A. R. and Hall, I. M. (2010). BEDTools: a flexible suite of utilities for comparing genomic features. Bioinforma. Oxf. Engl. 26, 841–842.

Raudvere, U., Kolberg, L., Kuzmin, I., Arak, T., Adler, P., Peterson, H. and Vilo, J. (2019). g:Profiler: a web server for functional enrichment analysis and conversions of gene lists (2019 update). Nucleic Acids Res. 47, W191–W198.

Righelli, D., Koberstein, J., Zhang, N., Angelini, C., Peixoto, L. and Risso, D. (2018). Differential Enriched Scan 2 (DEScan2): a fast pipeline for broad peak analysis. PeerJ Inc.

Russo, F., Righelli, D. and Angelini, C. (2016). Advancements in RNASeqGUI towards a Reproducible Analysis of RNA-Seq Experiments. BioMed Res. Int. 2016, 7972351.

Sakurai, H., Era, T., Jakt, L. M., Okada, M., Nakai, S., Nishikawa, S. and Nishikawa, S. (2006). In vitro modeling of paraxial and lateral mesoderm differentiation reveals early reversibility. Stem Cells Dayt. Ohio 24, 575–586.

Shen, L., Shao, N., Liu, X. and Nestler, E. (2014). ngs.plot: Quick mining and visualization of next-generation sequencing data by integrating genomic databases. BMC Genomics 15, 284.

Stoller, J. Z., Huang, L., Tan, C. C., Huang, F., Zhou, D. D., Yang, J., Gelb, B. D. and Epstein, J. A. (2010). Ash2l interacts with Tbx1 and is required during early embryogenesis. Exp Biol Med Maywood 235, 569–76.

Tanaka, M., Jokubaitis, V., Wood, C., Wang, Y., Brouard, N., Pera, M., Hearn, M., Simmons, P. and Nakayama, N. (2009). BMP inhibition stimulates WNT-dependent generation of chondrogenic mesoderm from embryonic stem cells. Stem Cell Res. 3, 126–141.

Turner, F. S. (2014). Assessment of insert sizes and adapter content in fastq data from NexteraXT libraries. Front. Genet. 5, 5.

Xu, H., Cerrato, F. and Baldini, A. (2005). Timed mutation and cell-fate mapping reveal reiterated roles of Tbx1 during embryogenesis, and a crucial function during segmentation of the pharyngeal system via regulation of endoderm expansion. Development 132, 4387–95.

Yan, J., Chen, S.-A. A., Local, A., Liu, T., Qiu, Y., Dorighi, K. M., Preissl, S., Rivera, C. M., Wang, C., Ye, Z., et al. (2018). Histone H3 lysine 4 monomethylation modulates long-range chromatin interactions at enhancers. Cell Res. 28, 204–220.

Yu, G., Wang, L.-G. and He, Q.-Y. (2015). ChIPseeker: an R/Bioconductor package for ChIP peak annotation, comparison and visualization. Bioinforma. Oxf. Engl. 31, 2382–2383.

Zang, C., Schones, D. E., Zeng, C., Cui, K., Zhao, K. and Peng, W. (2009). A clustering approach for identification of enriched domains from histone modification ChIP-Seq data. Bioinforma. Oxf. Engl. 25, 1952–1958.

