## Supplementary Figure 1 for "Chromatin and transcriptional response to loss of TBX1 in early differentiation of mouse cells"

# WT

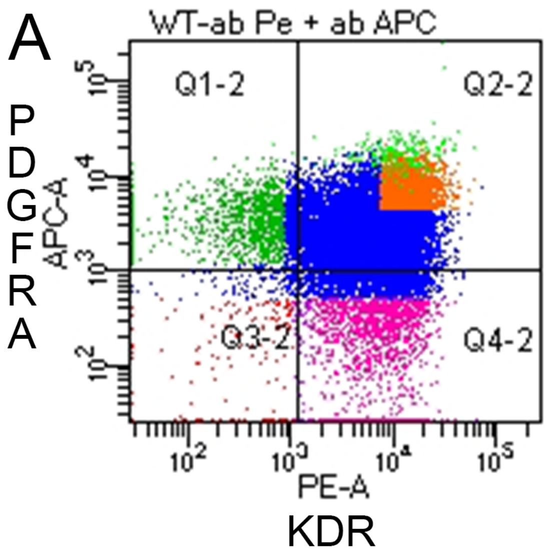


Q1-2= PDGFRα+/KDR- **9%**
Q2-2= PDGFRα+/KDR+ **68,2%**
Q3-2= PDGFRα-/KDR- **1,9%**
Q4-2= PDGFRα-/KDR+ **21%**

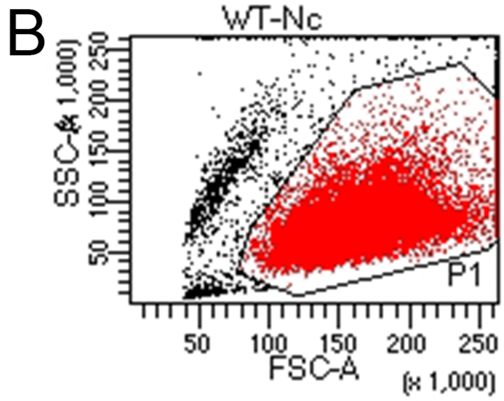


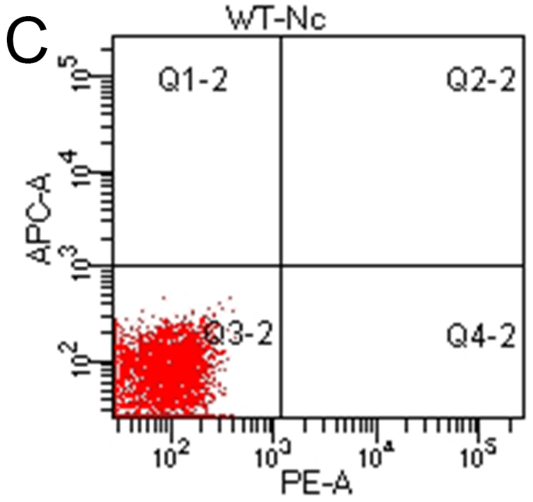


### *Tbx1-/-*

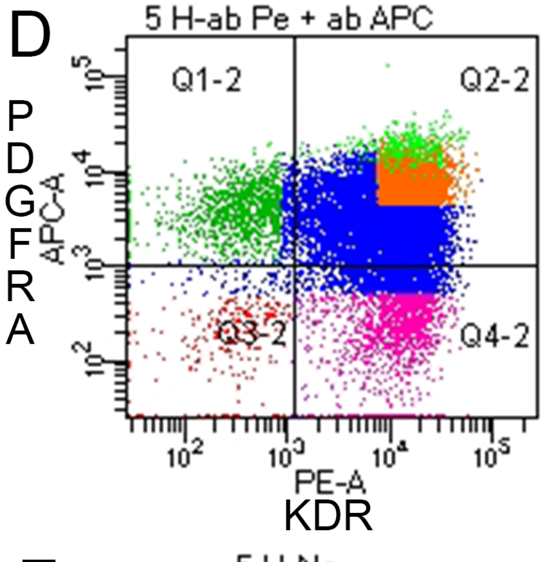


Q1-2= PDGFRα+/KDR- **8,3%**
Q2-2= PDGFRα+/KDR+ **71,8%**
Q3-2= PDGFRα-/KDR- **2,2%**
Q4-2= PDGFRα-/KDR+ **17,6%**

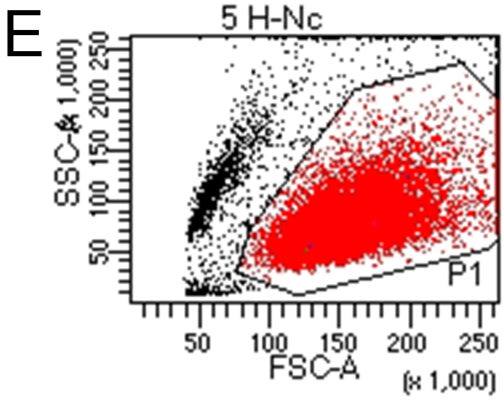


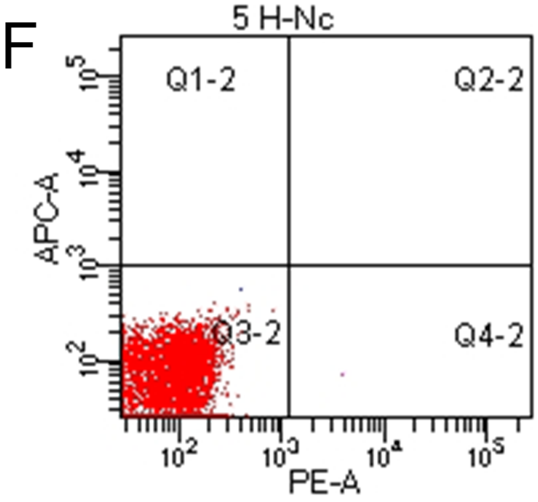
