## Supplementary figures and images for "Chromatin and transcriptional response to loss of TBX1 in early differentiation of mouse cells"

### Supplementary Figure 2

Supplementary Figure 2

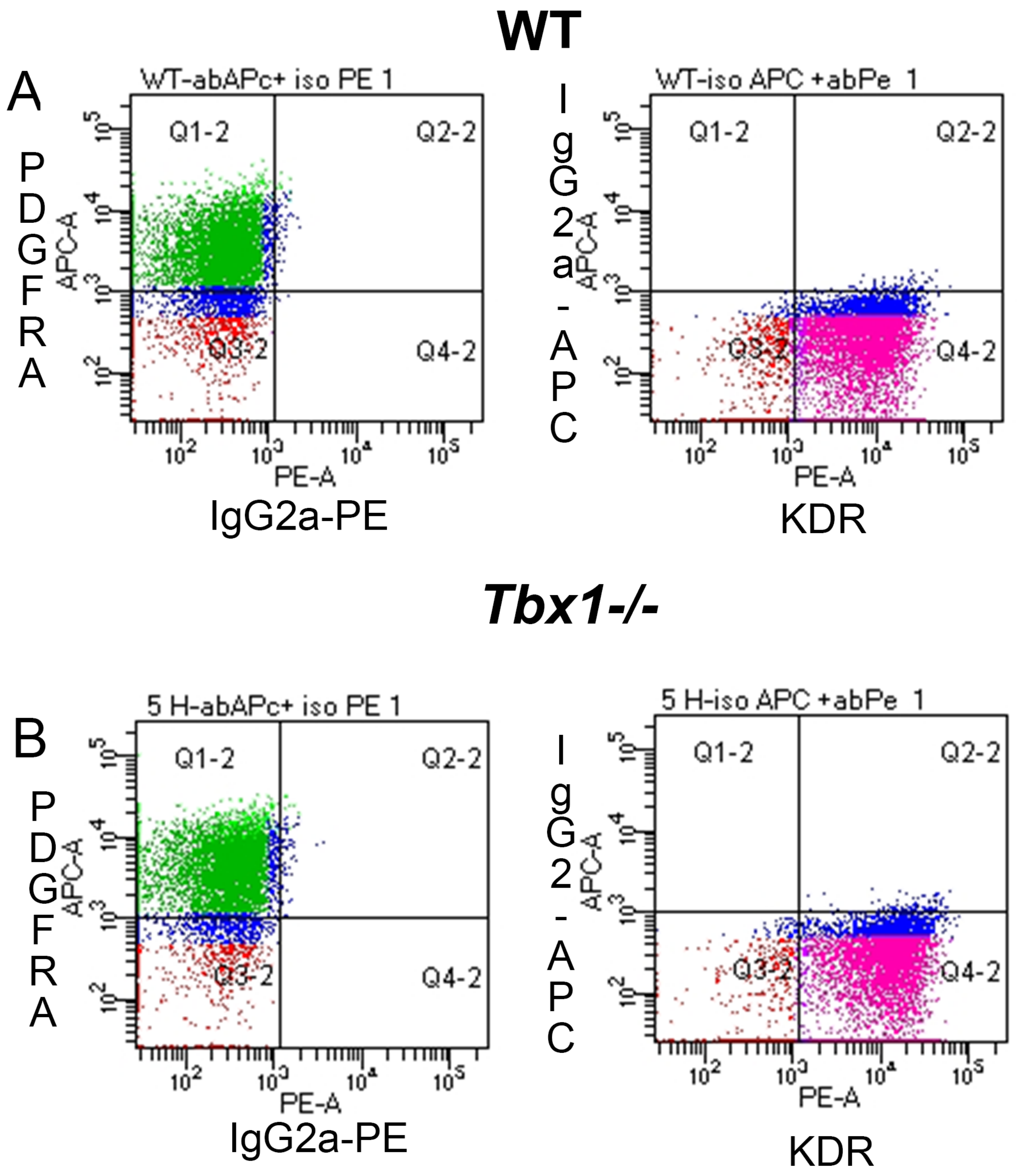
